# Whole-brain, gray and white matter time-locked functional signal changes with simple tasks and model-free analysis

**DOI:** 10.1101/2023.02.14.528557

**Authors:** Kurt G Schilling, Muwei Li, Francois Rheault, Yurui Gao, Leon Cai, Yu Zhao, Lyuan Xu, Zhaohua Ding, Adam W Anderson, Bennett A Landman, John C Gore

## Abstract

Recent studies have revealed the production of time-locked blood oxygenation-level dependent (BOLD) functional MRI (fMRI) signals throughout the entire brain in response to a task, challenging the idea of sparse and localized brain functions, and highlighting the pervasiveness of potential false negative fMRI findings. In these studies, ‘whole-brain’ refers to gray matter regions only, which is the only tissue traditionally studied with fMRI. However, recent reports have also demonstrated reliable detection and analyses of BOLD signals in white matter which have been largely ignored in previous reports. Here, using model-free analysis and simple tasks, we investigate BOLD signal changes in both white and gray matters. We aimed to evaluate whether white matter also displays time-locked BOLD signals across all structural pathways in response to a stimulus. We find that both white and gray matter show time-locked activations across the whole-brain, with a majority of both tissue types showing statistically significant signal changes for all task stimuli investigated. We observed a wide range of signal responses to tasks, with different regions showing very different BOLD signal changes to the same task. Moreover, we find that each region may display different BOLD responses to different stimuli. Overall, we present compelling evidence that the whole brain, including both white and gray matter, show time-locked activation to multiple stimuli, not only challenging the idea of sparse functional localization, but also the prevailing wisdom of treating white matter BOLD signals as artefacts to be removed.

## Introduction

Functional magnetic resonance imaging (fMRI) based on blood oxygenation-level dependent (BOLD) contrast is well-established for mapping cortical activity in the brain. By characterizing BOLD signal changes in response to known events or stimuli, task-based fMRI has proven successful at identifying and localizing brain regions that are functionally involved in response to these tasks or stimulations. Despite the common notion that sparse, specific brain regions are associated with specific tasks, evidence has also emerged that whole-brain hemodynamic changes may be evoked by brain neuronal responses to stimuli. Specifically, by using extensive fMRI acquisitions, resulting in high signal-to-noise ratio data, Gonzalez-Castillo et al. (1) showed that BOLD signal changes correlate with task timings throughout a majority of the cortex, with different BOLD responses in different regions of the grey matter in the brain. They demonstrate that the sparse activations observed in traditional fMRI maps result from elevated noise, and overly strict assumptions of BOLD response functions, in agreement with (and validated by) other studies showing increased activation with signal averaging (2-6), or regional differences in BOLD responses (3, 6-10). Thus, there is strong evidence for “whole-brain, time-locked activation” that can be revealed through signal averaging and model-free analysis.

However, “whole-brain” in this context refers to only the gray matter of the brain, whereas white matter BOLD signals are rarely reported, and in fact are often regressed out as nuisance covariates. However, multiple reports have demonstrated successful detection and analysis of BOLD signals in white matter (11, 12). For example, previous studies have reported white matter BOLD signal responses to tasks and at rest (13, 14), the relationships between white matter signals and the gray matter regions to which they connect (15, 16), and also robust, reproducible network properties of white matter BOLD signals (16, 17). Importantly, recent reports have confirmed that white matter responses to stimuli are similar to, but different from, those in gray matter (13, 14), generally indicating a slower response, smaller percent signal change, and variation across regions (18).

By analogy to the gray matter, it is unknown whether white matter displays a time-locked variation of BOLD signals across all structural pathways, and whether such changes in BOLD signals vary based on stimuli. We hypothesize the production of widespread white matter BOLD signal changes, even in pathways that are not specifically associated with a given task. To evaluate this, we averaged multiple trials of sustained stimuli to generate data with high signal-to-noise ratio, and use model-free analysis to assess signal changes in both gray matter regions and diffusion tractography-derived white matter pathways. We test whether significant signal changes are found throughout the white and gray matter, and additionally assess differences in BOLD signal changes across regions and across different stimuli.

## Methods

### Image acquisition

All imaging (**Figure 1, DATA**) was performed on a 3-T Philips Achieva CRX scanner (Philips Healthcare) at Vanderbilt University Institute of Imaging Science, using a 32-channel head coil. To provide anatomical references, high-resolution T1-weighted images were also acquired using a multishot, 3D gradient echo sequence at voxel size of 1 × 1 × 1 mm^3^. Next, for white matter pathway delineation, diffusion MRI images were acquired using a pulsed gradient spin echo (PGSE) echo planar imaging (EPI) sequence at a voxel size of 2 × 2 × 2 mm^3^, with 32 diffusion weighted images at a b-value of 1000 s/mm^2^, and a single non-diffusion weighted image.

**Figure 1.**
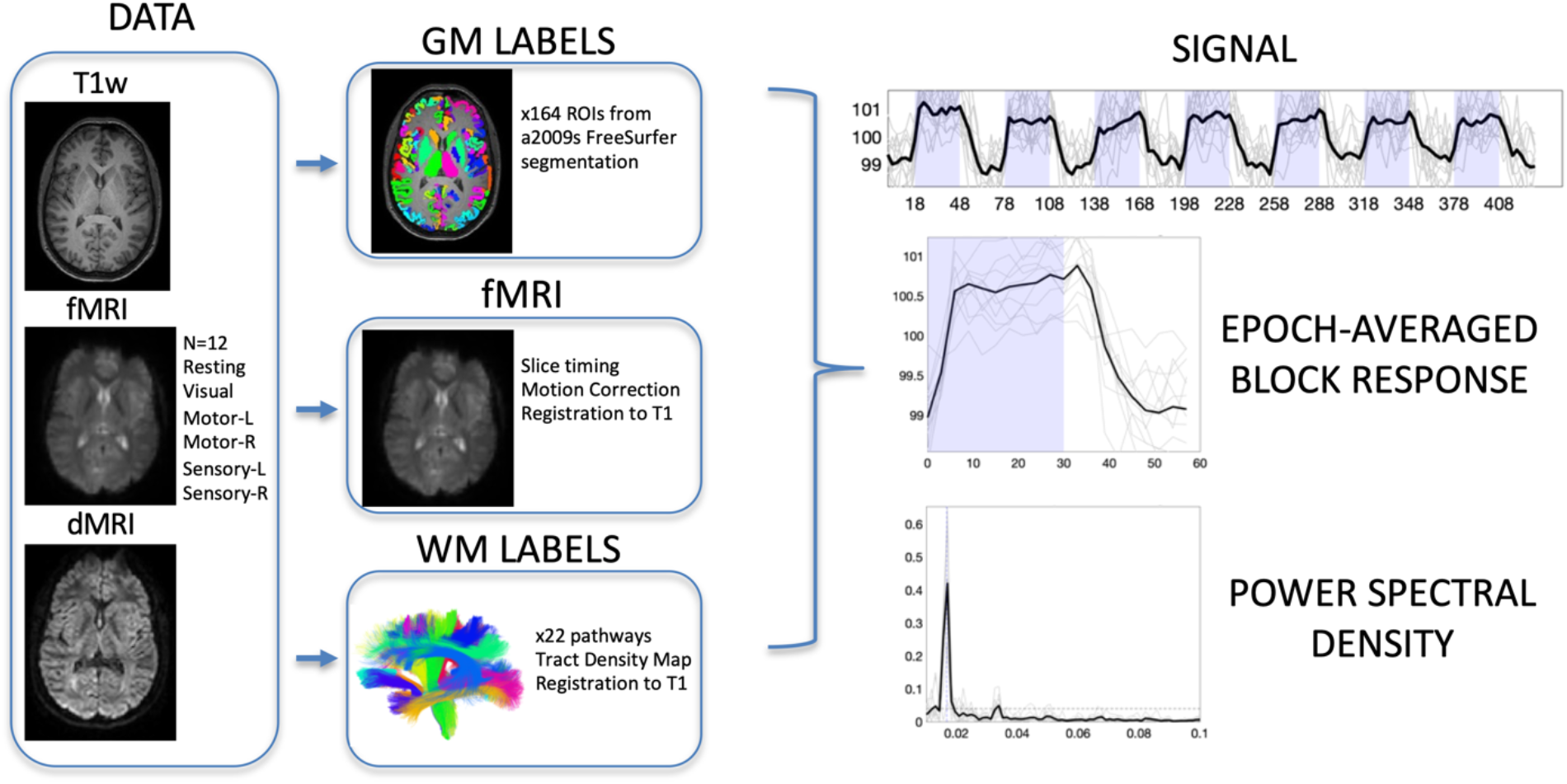
Image acquisition, preprocessing, region of interest delineation, and time-locked activation. Data includes T1-weighted structural images, fMRI data including resting state and 5 stimuli, and diffusion MRI. Gray matter labels were extracted from structural images, while white matter labels were derived from diffusion MRI fiber tractography. Functional MRI data were corrected for slice-timing and motion, and all data were registered to T1 space for analysis. Analysis included extracting region-averaged signal, from which the epoch-averaged block response can be visualized. Time-locked activation was determined by the power spectral density at task frequency (0.0167 Hz) with statistically significant activation determined by bootstrap analysis at p<0.01 with FDR correction.

Finally, six sets of images sensitive to BOLD contrast were acquired from these subjects using a single shot T2*- weighted gradient echo, echo planar imaging sequence with repetition time of 3s, echo time of 45 ms, SENSE factor of 2, matrix size of 80 × 80, field of view of 240 × 240 mm2, 43 axial slices of 3 mm thick with zero gap, and 145 sequential volumes. The BOLD images contained a resting state acquisition (Resting), and sets of images from sensory stimulations to the right hand and the left hand (Sensory-R and Sensory-L), motor tasks by the subjects’ right hand and left hand (Motor-R and Motor-L), and visual stimulations (Visual). All six runs had the same time duration of 435 s. During the image acquisitions, subjects lay in a supine position with eyes closed in a resting state or fixed on a screen mounted at the end of the scanner bore, viewed through mirrors mounted on the head coil. Head motions during image acquisitions were reduced by placing restricting pads within the head coil.

All stimulations and tasks were prescribed in a block design format, with five additional volumes acquired without stimulations or tasks at the beginning of each run. Sensory stimulations started with 30 s of hand stimulations by repeatedly brushing the palm followed by 30 s of no stimulations. The subjects fixed on an arrow sign pointing to the side of hand stimulation or a cross sign in the middle of the screen when there were no stimulations. Motor tasks started with 30 s of finger movement, in which subjects successively tapped each finger of the hand indicated by the arrow, followed by 30 s of no finger movements. In visual stimulations, the subjects continuously viewed the projection screen, which displayed 30 s of a checkerboard flashing at 8 Hz followed by 30 s of a cross sign on black ground. The blocks of stimulation or task were repeated seven times for each of the functional acquisitions. The total acquisition time was 435s, yielding 145 volumes for each task/stimulation.

In total, 15 subjects were included in this study (7 Male, 8 Female, age range 23-37), although not all BOLD contrasts were acquired on all subjects. Each BOLD contrast was represented N=12 times in our cohort.

### Image preprocessing

BOLD images were preprocessed (**Figure 1, fMRI**) using the statistical parametric mapping software package SPM12 (www.fil.ion.ucl.ac.uk/spm/software). The preprocessing performed corrections only for slice timing and head motions, with no subjects indicating large head motion (>2 mm in translation or >2° in rotation). Importantly, no spatial smoothing was applied to any fMRI data so as to minimize partial volume effects. The diffusion data were preprocessed to correct for motion and eddy currents using the FSL software package (19).

### Region of Interest delineation

Subject specific labels for gray and white matter were created. FreeSurfer (20) was run on all T1 weighted images, and gray matter labels were extracted from the Destrieux (21) atlas parcellation, resulting in 164 gray matter labels (**Figure 1, GM LABELS**) already in the space defined by the anatomical T1-weighted image.

Fiber tractography was performed on the diffusion MRI data using the TractSeg tool (22) that performed probabilistic bundle-specific tractography to generate streamlines representing the spatial trajectories of white matter pathways. Twenty two major association, projection, and commissural pathways were selected for this study (**Figure 1, WM LABELS**), and tract density images were generated that described the density of streamlines in each imaging voxel. Diffusion data were registered with the T1-weighted image using FSL’s epi_reg tool for EPI to structural registration, and white matter labels were transformed to the structural image space.

Finally, the mean FMRI image was registered to the T1-weighted image using FSL’s epi_reg tool. BOLD signal, for all contrasts, was transformed to the structural image space. At this point, white and gray matter labels, and fMRI signal are all aligned within the same space.

### Time-locked activation

Our aim was to determine which regions of interest indicated time-locked activation with the stimulus/task across a population. To do this, we first extracted the averaged fMRI signal within each of the white and gray matter regions of interest (**Figure 1, SIGNAL**). From this, the power spectral density was estimated via the periodogram method (periodogram function in MatLab) which represented the distribution of power per unit frequency (**Figure 1, POWER SPECTRAL DENSITY**). In this study, the frequency of interest occurred at 0.0167Hz (1/60s). To determine whether there was significant time locked activation (i.e., statistically significant power at this frequency), we performed a bootstrap analysis of the resting state data to generate our null distribution of power at this frequency. Here 10,000 bootstrap signals were generated from volumes of 39×39×39mm3 (13×13×13 voxels) within 12 subjects, and the average power spectral density at 0.0167Hz was calculated. From this, given the real data (i.e., the subject averaged power spectral density in a region of interest), we generated a p-value for statistically significant time-locked activation. Regions with p<0.01 after false discovery rate correction are considered time-locked, or activated, by our tasks, and the epoch-averaged response function (**Figure 1, EPOCH-AVERAGED BLOCK RESPONSE**) can be visualized.

## Results

### Widespread Gray Matter Time-locked Activation

**Figure 2** shows the subject-averaged signal and epoch-averaged responses for several selected gray matter regions for the visual task. Many regions show both qualitatively observable, and statistically significant, time-locked activation with the task. While some regions show the canonical sustained boxcar response, or a sustained response with small transitory response at the onset or offset of the task, other trends are also apparent, including delayed responses, continuously changing signals, or multiple peaks. The observation of widespread gray matter time-locked activation is in agreement with previous literature (1).

**Figure 2.**
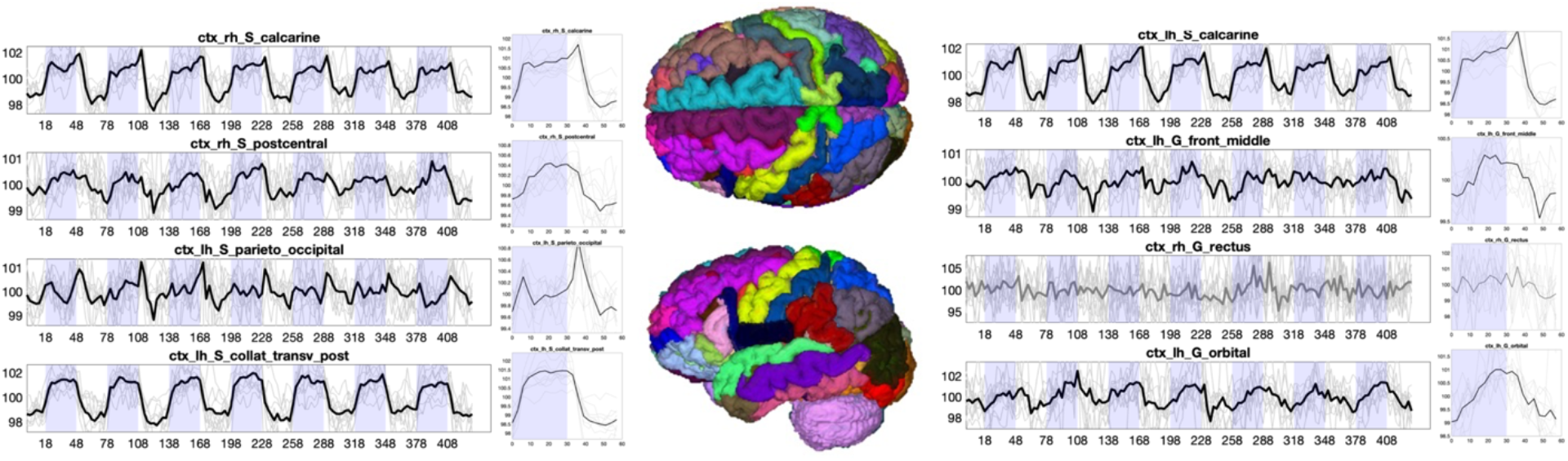
Time-locked activation in gray matter. The signal and epoch-averaged response are shown for 8 selected gray matter regions. Plots are shown in BOLD if statistically significant time-locked activation is present, and mean value is shown as a thick line, with data from all subjects (N=12) as thin lines.

### Widespread White Matter Time-locked Signal Changes

Beyond the traditional gray matter analysis, we show white matter signals for several selected pathways in **Figure 3**. White matter also shows widespread time-locked signal changes across projection, commissural, and association pathways. Here, several pathways show strong, canonical boxcar shapes, while others have a somewhat smaller signal change, or different shapes including, for example, negative signal changes during stimulation.

**Figure 3.**
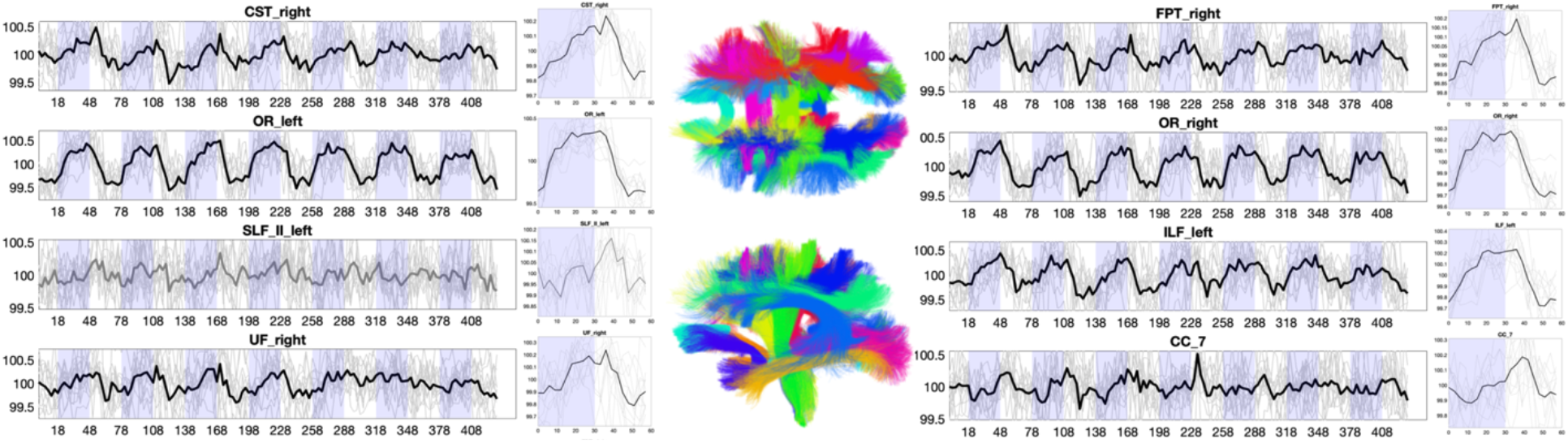
Time-locked BOLD signal changes in white matter. The signal and epoch-averaged responses are shown for 8 selected white matter regions. Plots are shown in BOLD if statistically significant time-locked signal changes are present, and mean value is shown as a thick line, with data from all subjects (N=12) as thin lines.

### Variation in tasks

**Figure 4** shows the signal for two selected gray matter and two selected white matter regions across all BOLD contrasts/tasks, including resting state. First, resting state does not show time-locked activation at 0.0167Hz. However, all other tasks, for both gray matter and both white matter regions, show statistically-significant signal changes. Notably, the signal response for all regions varies dramatically based on the tasks, or hemisphere in which the task is performed.

**Figure 4.**
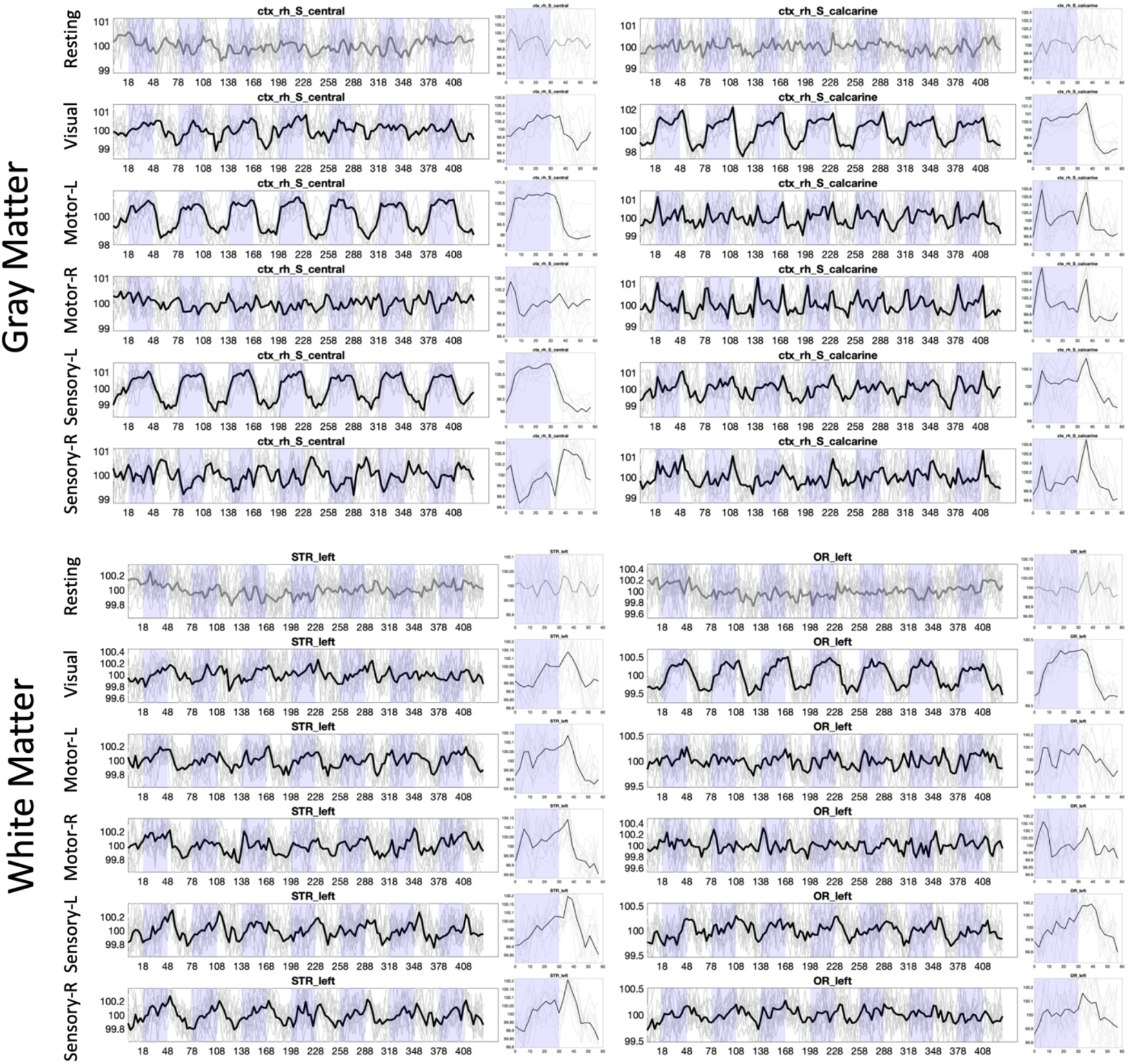
Time-locked activation varies across regions and across stimuli, in both white and gray matter. Example signal and epoch-averaged signal are shown for two gray matter and two white matter regions, across all tasks. Plots are shown in BOLD if statistically significant time-locked activation is present.

### Whole-brain time-locked activation

**Figure 5** visualizes regions based on the power spectral density (showing only statistically significant regions) for all tasks. Here, the whole-brain response, for all tasks, is clearly visible, importantly in both gray and white matters. The traditional regions of interest associated with each function indeed show the strongest activations, for example occipital lobe for visual tasks or pre/post-central sulci for motor tasks. Additionally, so does the white matter directly projecting to/from these regions, the optic radiations in visual tasks, the corticospinal tract for motor/sensory tasks. However, for all tissues nearly all regions show significant signal changes, including those that play accessory roles in tasks or are not hypothesized to play any role at all.

**Figure 5.**
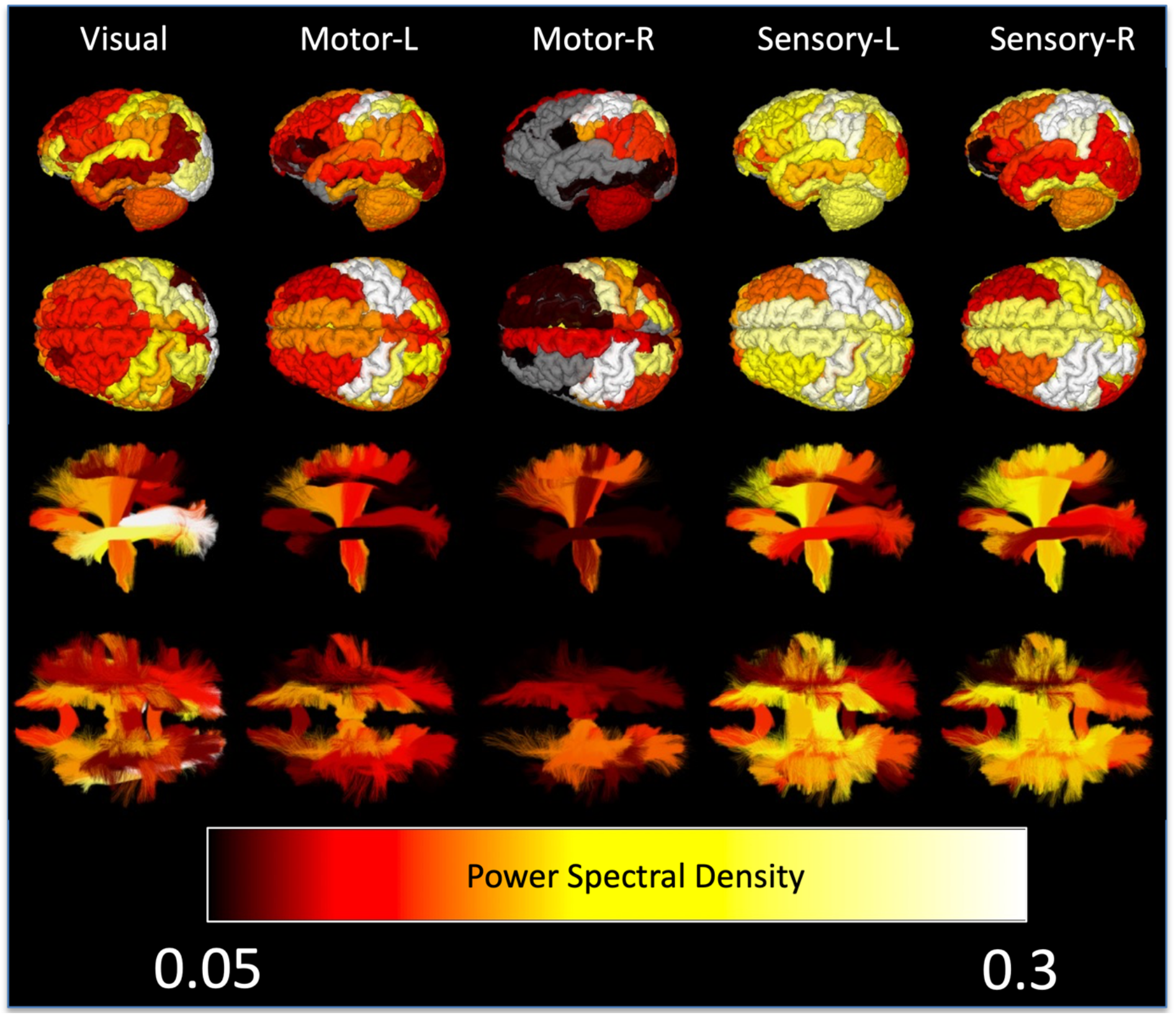
Whole brain time locked signal changes in both white and gray matter. Gray matter regions (top) and white matter pathways (bottom) are shown color-coded based on subject-averaged power-spectral density for 5 tasks. Non-significant changes are shown in gray for gray matter, and not visualized in white matter.

## Discussion

### White Matter BOLD Signal

Using simple tasks and model-free analysis, we show here that task-locked BOLD signal changes may be evoked across the whole brain, in both white and gray matter. BOLD signal in white matter remains controversial (11, 12), as the blood flow and volume in white matter are much lower than those in the gray matter (23, 24), and the biological basis of BOLD signal mechanisms in white matter is unclear. While there is clear evidence that the fMRI signal in gray matter corresponds to changes in local post-synaptic potentials (25), the evidence for BOLD signal fluctuations caused by propagation of action potentials is less clear (12), though white matter also contains a large number of glial cells which are engaged in neural activity and consume energy. However, reports of white matter fMRI signal changes are becoming more frequent, with studies commonly describing activations in the corpus callosum (26-29), internal capsule (30, 31), and optic radiations, among other large white matter pathways (13, 32). Here, we confirm detectable signal fluctuations in a large number of projection, commissural, and association pathways.

Gawryluk et al. (12) and Gore et al. (2019) describe several explanations for why white matter BOLD signals may have not been detected in previous studies, or are dismissed entirely as artifacts. First, white matter BOLD signals and signal changes are lower in magnitude than gray matter activation (12) as expected by reason of lower blood volume. We therefore performed multiple within-subject and across-subject averaging to improve SNR. Moreover, in conventional analyses, clustering of voxels to increase SNR usually is performed over cubic or spherical volumes that are unsuited to thin white matter tracts. Here we average the signal over the entire length of each white matter pathway. Second, fMRI analysis has been tailored towards gray matter data. For example, white matter signals may be treated as nuisance regressors (33, 34), or used as a “zero activity” references, and a specific form has been assumed for the hemodynamic response function in General Linear Models which may not be appropriate for white matter (35) (see below for more discussion). Here, we used model-free analysis, making no assumptions about the shape of the BOLD signal changes.

BOLD signal changes in white matter could also be attributed to task-related motion (12). However, voxels near the boundaries of tissue would be most susceptible to this artifact whereas we have weighted the white matter signal by density within the white matter, and there is no obvious reason why white matter would be any more susceptible to this than gray matter. Finally, white matter activation could be attributed to partial volume effects from gray matter, especially if using larger voxels in image acquisitions, or after smoothing data. Here, to minimize partial volume effects, we did not smooth the data. Moreover, by weighting by tract density, the white matter signal was heavily weighted by the dense core of deep white matter bundles for the large pathways studied here (Figure 6), reducing potential gray matter confounds.

**Figure 6.**
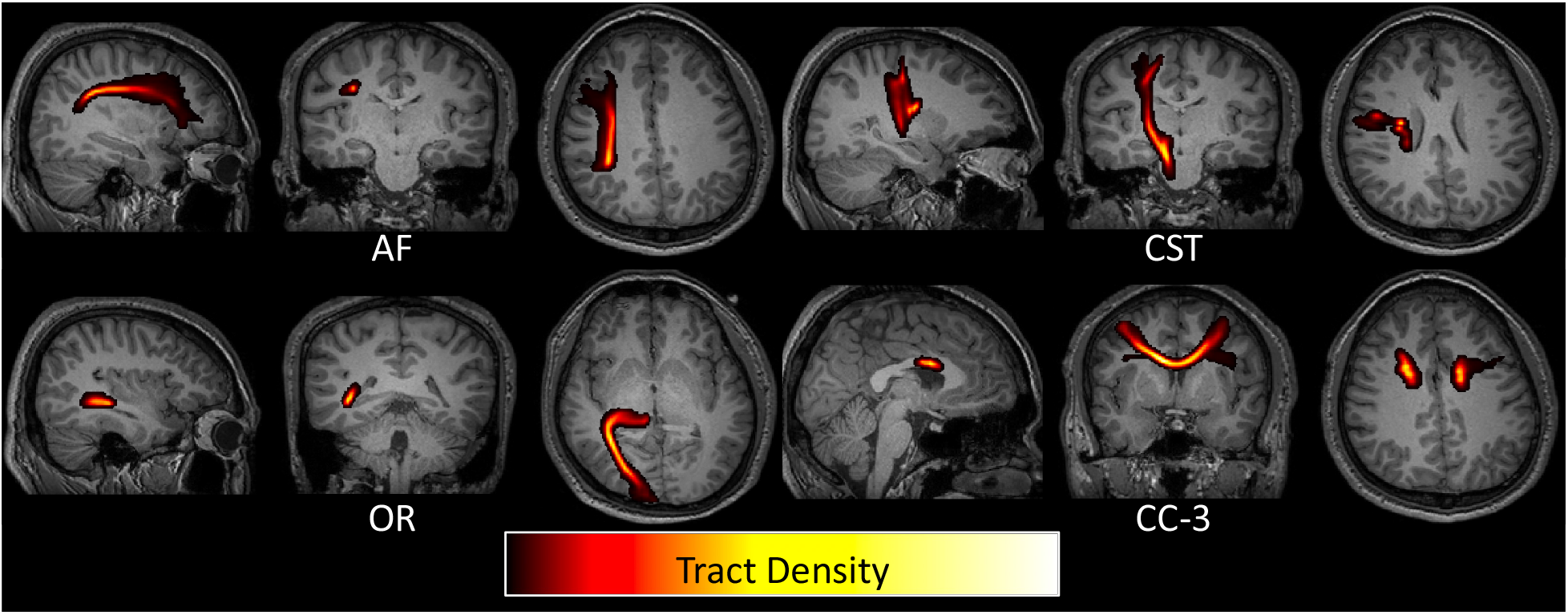
Weighting white matter BOLD signal by tract density reduces partial volume effects with gray matter regions. Four example pathways are shown as tract density maps overlaid on T1 images, where the highest density of streamlines is consistently in deep white matter regions.

We provide strong evidence that evoked white matter BOLD signals are near ubiquitous within the brain when a task or stimulus is presented, and that they occur concurrently with widespread activity in grey matter as has been previously reported. Their origins remain to be more clearly established, but they share multiple features with the hemodynamic responses that are attributed to neural activity in gray matter. Signal propagation along axons reportedly requires only a small fraction of the total white matter energy budget (23, 24), but other processes within white matter, including those in other cell types, may provoke a BOLD response to functional activity in the brain.

### Brain Wide Activation

Our study was motivated in part by findings from (1), which found non-random gray matter-wide modulations of BOLD responses to tasks. These results raise the issue that many fMRI studies may overlook involvement of many brain areas due to insufficient power to detect activations. Statistical parametric maps, which describe involvement of brain regions in a task, are usually derived by fitting empirical BOLD signal data to a model of the BOLD response function and evaluating quality of fit. As highlighted throughout the literature, inappropriate models of the hemodynamic response function may decrease statistical power to detect true effects, resulting in an underestimation of the amount of variance explained by the signal (and overestimation of noise) and substantial number of false negative results (35).

Even for a simple task such as a visual discrimination task (1), almost the entire brain shows BOLD signal changes, and the act of labeling brain areas as active versus inactive is likely overly ambiguous. Here, we extend this work, showing that in addition to visual stimulation, also sensory and motor tasks produce wide-spread signal changes outside of the traditional locations associated with these tasks. These widespread effects may reflect neural engagement of multiple areas or some form of vascular coupling between regions. Intuitively, if the entire cortex is experiencing changes, the white matter substrates linking these cortical regions must also be consequently affected (or effecting these changes). Regardless of the biological basis of the signal in WM, it is clear that some change in blood volume, oxygenation, or relaxation rate occurs in a large portion of the white matter pathways of the brain. These white matter pathways also do not have single-task functions, but show BOLD signal changes in a multitude of tasks. However, here, there is significant spatial overlap of structural pathways within the deep white matter (36, 37) which may confound the identity of the origins of some white matter signals.

### Linking Structure and Function

One goal of brain mapping by MRI is to establish links between neuronal substrates and their structural connections in order to better understand their functional relevance. Structure and function in neuroimaging have traditionally been studied through independent measures. While functional engagement has traditionally been identified through BOLD signals and focused on gray matter, structural connections have been quantified through diffusion MRI and focused on white matter. We propose that both BOLD and diffusion contrasts can provide useful information on structural and functional characteristics of both tissue types. Here, fMRI shows robust signal changes in white matter regions. Further studies may elucidate relationships between white and gray matter signals, including potential time delays and differences in stimuli. It may be possible, using high temporal resolution and high signal to noise ratio data, to identify which white matter voxels are associated with connections to specific cortical regions, providing validation for diffusion tractography, which also suffers from both false positive and false negative detections (38-41). Further, studies have shown that BOLD signals in white matter pathways are highly correlated to other voxels within the same pathway (42), offering a way to either parcellate white matter based on functional signals or perform fiber tractography directly on functional data (43).

Diffusion MRI has also been claimed to detect functional activations as changes in apparent diffusion coefficients (44). Task-correlated changes in diffusivity have been hypothesized to represent structural changes such as cell-swelling during neuronal activation (44), and have shown activation in similar regions to that of traditional BOLD contrast (45, 46). These types of diffusion experiments can be utilized in the white matter (47-49), and future investigations may enable teasing out microstructural contributions to the white matter signal described in the current study.

## Acknowledgements

This work was supported by the National Science Foundation Career Award #1452485, the National Institutes of Health under award numbers R01EB017230, K01EB032898, and in part by ViSE/VICTR VR3029 and the National Center for Research Resources, Grant UL1 RR024975–01.

